# A Suite of Eight *Toxoplasma gondii* Effectors Cooperates to Activate the Non-canonical NF-κB Pathway

**DOI:** 10.64898/2026.03.12.711255

**Authors:** Kenna Berg, Michael Panas, Samarchith P Kurup, John C Boothroyd, Alex Rosenberg

**Affiliations:** Department of Infectious Diseases, University of Georgia, Athens, GA, USA; Department of Cellular Biology, University of Georgia, Athens, GA, USA; Center for Tropical and Emerging Global Diseases, University of Georgia, Athens, GA, USA; Department of Microbiology and Immunology, Stanford University School of Medicine, 299 Campus Drive, Stanford, CA 94305, United States

**Keywords:** Toxoplasma gondii, non-canonical NF-κB, dense granule effectors, MYR translocon, RelB, p52, NIK stabilization, TRAF3

## Abstract

As a master of host-cell reprogramming, *Toxoplasma gondii* (*T. gondii*) tachyzoites manipulate diverse signaling networks to establish a niche permissive for long-term infection. While the parasite’s subversion of canonical NF-κB signaling (p65/p50) is well established, how infection impacts the non-canonical NF-κB pathway has been largely unexplored. Here, we report that *T. gondii* infection induces robust nuclear accumulation of the non-canonical NF-κB subunits RelB and p52 in both human and murine cells. This activation follows a gradual kinetic profile and is conserved across both Type I and Type II parasite genetic backgrounds.

We demonstrate that this reprogramming is strictly dependent on the MYR1-dependent export of dense granule effectors. Mechanistically, *T. gondii* infection drives the depletion of the negative regulator TRAF3, leading to the stabilization of NF-κB-inducing kinase (NIK), phosphorylation of p100, and its subsequent processing into p52. Utilizing a panel of combinatorial knockout parasites, we reveal that no single effector is responsible for this phenotype. Instead, a suite of eight MYR1-dependent effectors, IST, NSM, HCE1/TEEGR, GRA16, GRA18, GRA24, GRA28, and GRA84, functions through a collaborative, additive network to trigger the non-canonical response. These findings highlight a distributed regulatory strategy used by the parasite to overcome host transcriptional robustness and shape host signaling.

**Importance:** *Toxoplasma gondii* infects nearly one-third of the global population and establishes infection by extensively rewiring host immune signaling. While decades of work have focused on how the parasite modulates canonical NF-κB activity, whether it also engages the alternative, non-canonical arm of this pathway has remained unclear. Here, we show that *T. gondii* tachyzoites activate non-canonical NF-κB signaling, driving nuclear accumulation of RelB/p52 through MYR1-dependent effector export. Unexpectedly, no single effector is responsible. Instead, eight secreted proteins act cooperatively to enable NIK stabilization and engage the alternative NF-κB cascade, revealing a networked mode of immune control. This discovery highlights a regulatory logic evolved by the parasite to overcome host transcriptional robustness. Together, these findings identify non-canonical NF-κB activation as a new axis of host-parasite interaction and expand our understanding of how *T. gondii* reprograms central immune signaling circuits through multi-effector networks.

## Introduction

*Toxoplasma gondii* (*T. gondii*) is an obligate intracellular apicomplexan parasite that infects approximately one third of the human population (1). It causes severe disease in immunocompromised individuals and congenitally infected infants (2). In North America and Europe, the parasite population is dominated by a small number of clonal lineages: Type I strains are acutely virulent, while Type II and Type III strains show intermediate and low virulence in laboratory mice, respectively (3). A central feature of *T. gondii* pathogenesis is the establishment of the parasitophorous vacuole (PV), a specialized niche that protects the parasite while allowing it to reprogram host cell pathways (4). This remodeling is driven by the delivery of effector proteins from rhoptries and dense granules into the host cell (5).

The transport of exported dense granule effectors across the parasitophorous vacuole membrane (PVM) into the host cell appears to be mediated by a dedicated translocon complex composed of MYR1-MYR4 proteins (6–9). This machinery is supported by the ASP5 protease, GRA45 chaperone, ROP17 kinase, GRA44 phosphatase, and the accessory protein GRA59 (10–13). To date, eight MYR1-dependent exported dense granule effectors have been identified, the majority of which localize to the host nucleus: IST, NSM, HCE1/TEEGR, GRA16, GRA18, GRA24, GRA28, and the recently characterized host nuclear effector GRA84 (14–23). MYR1-dependent effectors account for the majority of *T. gondii-*driven host transcriptional reprogramming (24). Our recent work and that of others indicates that *T. gondii* effectors often function in redundant, synergistic, or antagonistic networks rather than as isolated factors. This combinatorial logic is prominent in the subversion of two major host pathways: IFN-γ and NF-κB (25, 26).

In the subversion of the IFN-γ signaling pathway, IST blocks STAT1-dependent transcription by recruiting Mi-2/NuRD to STAT1-bound promoters and preventing coactivator engagement, while NSM reinforces repression during chronic infection by elevating NCoR/SMRT repressor activity (14, 16). In parallel, functional genetic studies support a redundant network of four MYR-dependent nuclear effectors, IST, GRA16, GRA24, and GRA28, that collectively suppress IFN-γ-driven programs and prevent IFN-γ-associated host cell death and premature parasite egress (25).

A similar, combinatorial logic shapes NF-κB-mediated inflammatory signaling during infection. In Type II strains, GRA15 serves as a dominant driver of canonical NF-κB activation by recruiting TRAF2 and TRAF6 to promote IKK activation and p65 nuclear translocation, with additional effectors and feedback mechanisms shaping p65 nuclear accumulation and downstream cytokine output (27–29). GRA24 further tunes this response by sustaining nuclear accumulation of c-Rel (30). Other effectors restrain pathway output to limit pathology, including the nuclear repressor HCE1/TEEGR, which recruits EZH2-linked machinery to silence subsets of NF-κB target genes (18). In infected macrophages, these same NF-κB inputs are repurposed toward dissemination by promoting NF-κB-dependent CCR7 expression. GRA28 acts as a central enhancer by increasing chromatin accessibility at the Ccr7 locus via SWI/SNF, thereby potentiating GRA15/24-driven, NF-κB-dependent transcription. Together, these activities drive CCR7-dependent hypermigration, with additional support from GRA16 and GRA18 and counter-regulation by HCE1/TEEGR (26, 31).

Despite extensive attention to canonical NF-κB regulation during *T. gondii* infection, whether and how the parasite engages the alternative, non-canonical NF-κB pathway has remained less clear (32). The non-canonical pathway is defined by inducible processing of p100 into p52 and subsequent nuclear accumulation of RelB and p52 complexes, typically downstream of a subset of TNF-receptor family members and related signaling axes (33). Mechanistically, non-canonical activation is governed by the stability of NF-κB-inducing kinase (NIK) (34). Under resting conditions, NIK is kept low through a TRAF2, TRAF3, and cIAP-dependent ubiquitination system; pathway activation involves TRAF3 degradation and NIK stabilization, leading to IKKα-driven phosphorylation of p100 and its processing to p52 (35).

Early studies demonstrated that this pathway plays a critical role during *T. gondii* infection. RELB-deficient mice are highly susceptible to infection, succumbing rapidly due to a profound failure in the infection-induced activation of NK cells and the production of protective IFN-γ by T cells (36). While these foundational studies underscored the necessity of RelB for both innate and adaptive immunity, the molecular strategies employed by *T. gondii* to interact with this pathway have remained unknown.

Here, we identify activation of the non-canonical NF-κB pathway as a previously unrecognized consequence of *T. gondii* infection. Surprisingly, this response does not appear to be explained by any single exported effector but instead emerges from a distributed mode of host cell reprogramming. These findings reveal a new dimension of *Toxoplasma*-host interaction and suggest that parasite engagement of non-canonical NF-κB mirrors the multilayered regulatory logic that governs this pathway in uninfected cells.

## Results

### MYR1-dependent effector export drives nuclear accumulation of RelB and p52

MYR1-dependent export of dense granule effectors is a major driver of host-cell reprogramming, reshaping multiple core signaling pathways during infection (24). To identify dominant transcriptional regulators associated with this MYR1-dependent response, we performed *in silico* transcription-factor-enrichment analysis on host transcripts induced by WT parasites but not by the *Δmyr1* mutant. This analysis identified RelB as a top hit, implicating activation of the non-canonical NF-κB pathway (Fig. 1A).

**Figure 1.**
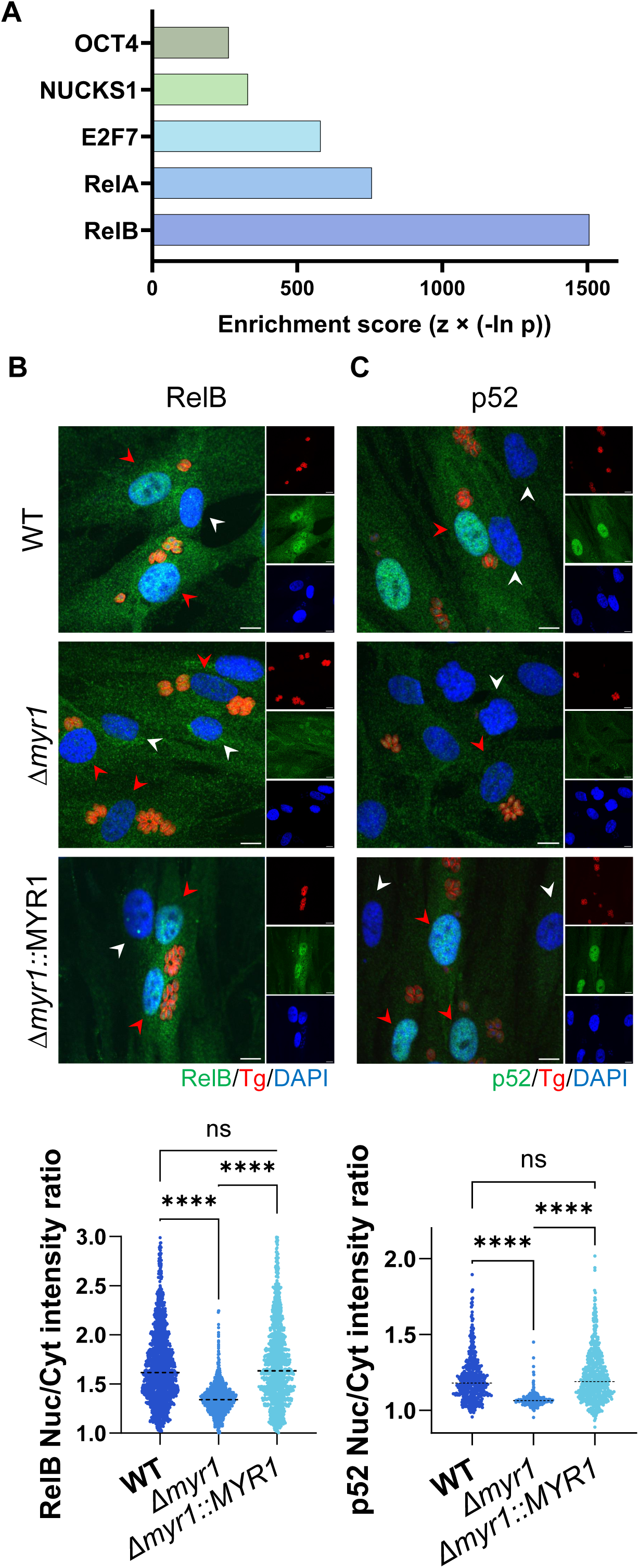
MYR1-dependent effector export drives nuclear accumulation of RelB and p52. **A)** Top-scoring transcription factors predicted to regulate the MYR1-dependent upregulated genes in HFF cells relative to uninfected controls were identified by analysis of previously published data (24) using Enrichr. **B-C)** Representative images (top) and quantitative analysis (bottom) for RelB **(B)** and p52 **(C)** nuclear accumulation in HFFs infected with RH (WT), Δ*myr1* or Δ*myr1*::MYR1 complement parasites. Twenty-four hours post-infection, cells were fixed and labeled with DAPI (blue), anti-GAP45 (red), and either anti-RelB or anti-p52 (green). Scale bars = 10 µm. Plots display the nuclear-to-cytoplasmic signal ratios for at least 300 infected cells (red arrows) per condition; uninfected cells are indicated by white arrows. Data from three independent experiments were combined for analysis. The horizontal dashed lines indicate the mean. Statistical significance was determined using a one-way ANOVA with Tukey’s multiple comparison test; ****P < 0.0001, ns = not significant.

Guided by this prediction, we examined RelB expression and localization in human foreskin fibroblasts (HFFs) infected with Type I (RH) or Type II (ME49) *T. gondii* strains. Immunofluorescence imaging combined with quantitative analysis revealed that the RelB nuclear-to-cytoplasmic ratio increased to ∼1.68 specifically in wild-type (WT) infected cells (Fig. 1B, Fig. S1A). This nuclear accumulation was abolished during infection with Δ*myr1* parasites and restored upon MYR1 complementation (Δ*myr1*::MYR1), indicating that MYR1-dependent effector export is likely required for RelB nuclear enrichment (Fig. 1B, Fig. S1A).

We next tested whether this phenotype is conserved across host species. Although uninfected mouse embryonic fibroblasts (MEFs) exhibited higher basal nuclear RelB than HFFs, infection with WT parasites of either Type I or Type II strains still induced a significant ∼1.3-fold increase in nuclear RelB (Fig. 2A and S2A). As in human cells, this increase was eliminated during Δ*myr1* knockout infection and regained upon complementation (Δ*myr1*::MYR1), supporting a conserved MYR1-dependent mechanism (Fig. 2A and S2A).

**Figure 2.**
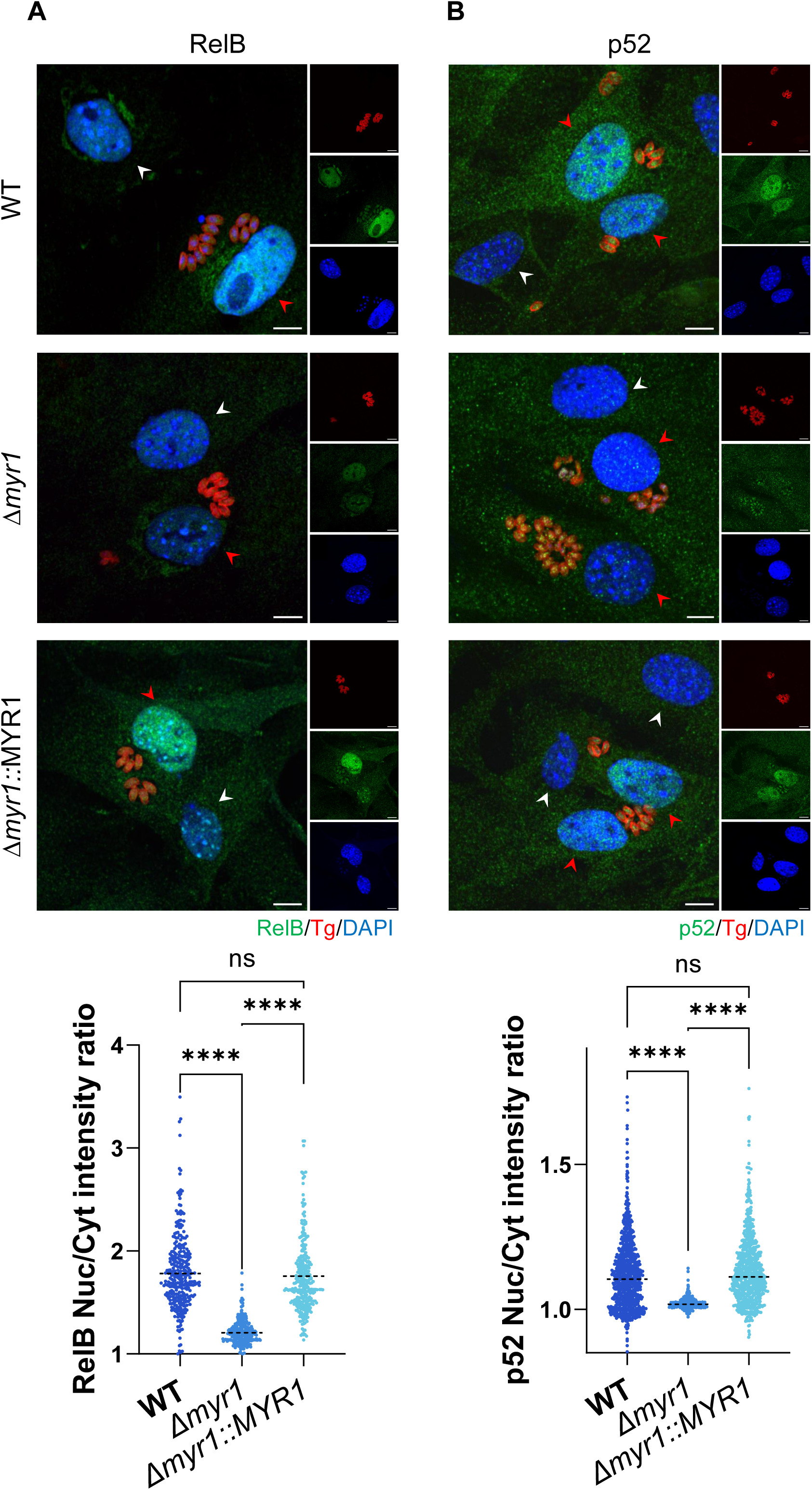
MYR1-dependent RelB and p52 nuclear accumulation in murine cells. **A-B)** Representative images (top) and quantitative analysis (bottom) for RelB **(A)** and p52 **(B)** nuclear accumulation in MEFs infected with RH (WT), Δ*myr1* or Δ*myr1*::MYR1 complement parasites. Twenty-four hours post-infection, cells were fixed and labeled with DAPI (blue), anti-GAP45 (red), and either anti-RelB or anti-p52 (green). Scale bars = 10 µm. Plots display the nuclear-to-cytoplasmic signal ratios for at least 300 infected cells (red arrows) per condition; uninfected cells are indicated by white arrows. Data from three independent experiments were combined for analysis. The horizontal dashed lines indicate the mean. Statistical significance was determined using a one-way ANOVA with Tukey’s multiple comparison test; ****P < 0.0001, ns = not significant.

Under basal conditions, RelB is sequestered in the cytoplasm by the NF-κB2 precursor p100, which functions as an IκB-like inhibitor through its C-terminal ankyrin repeat domains. Pathway activation triggers partial proteasomal processing of p100 into the active p52 subunit, facilitating the nuclear translocation of RelB/p52 heterodimers (37, 38). Consistent with this mechanism, infection with either Type I or Type II *T. gondii* strains increased nuclear p52 in both HFFs and MEFs (Fig. 1C, 2B, S1B, S2B). In all cases, nuclear p52 accumulation was lost during infection with *Δmyr1* parasites and restored by complementation (*Δmyr1*::MYR1) (Fig. 1C, 2B, S1B, S2B), indicating that MYR1-dependent effector export is apparently required for the processing event associated with RelB nuclear entry.

### MYR1-dependent TRAF3 depletion stabilizes NIK and drives p100 processing

The non-canonical NF-κB pathway depends on NIK stabilization and proteasome-dependent processing of p100, rather than rapid IκB degradation, and therefore typically displays slower, more sustained activation kinetics than the canonical pathway (33, 39). To test whether *T. gondii* infection follows similar dynamics, we quantified nuclear RelB levels in infected HFF cells every 6 hours over a 24-hour period. This analysis revealed a gradual, progressive increase in nuclear RelB that reached a maximum at 18 hours, consistent with the characteristic kinetics of the NF-κB non-canonical arm (Fig. 3A) (40).

**Figure 3.**
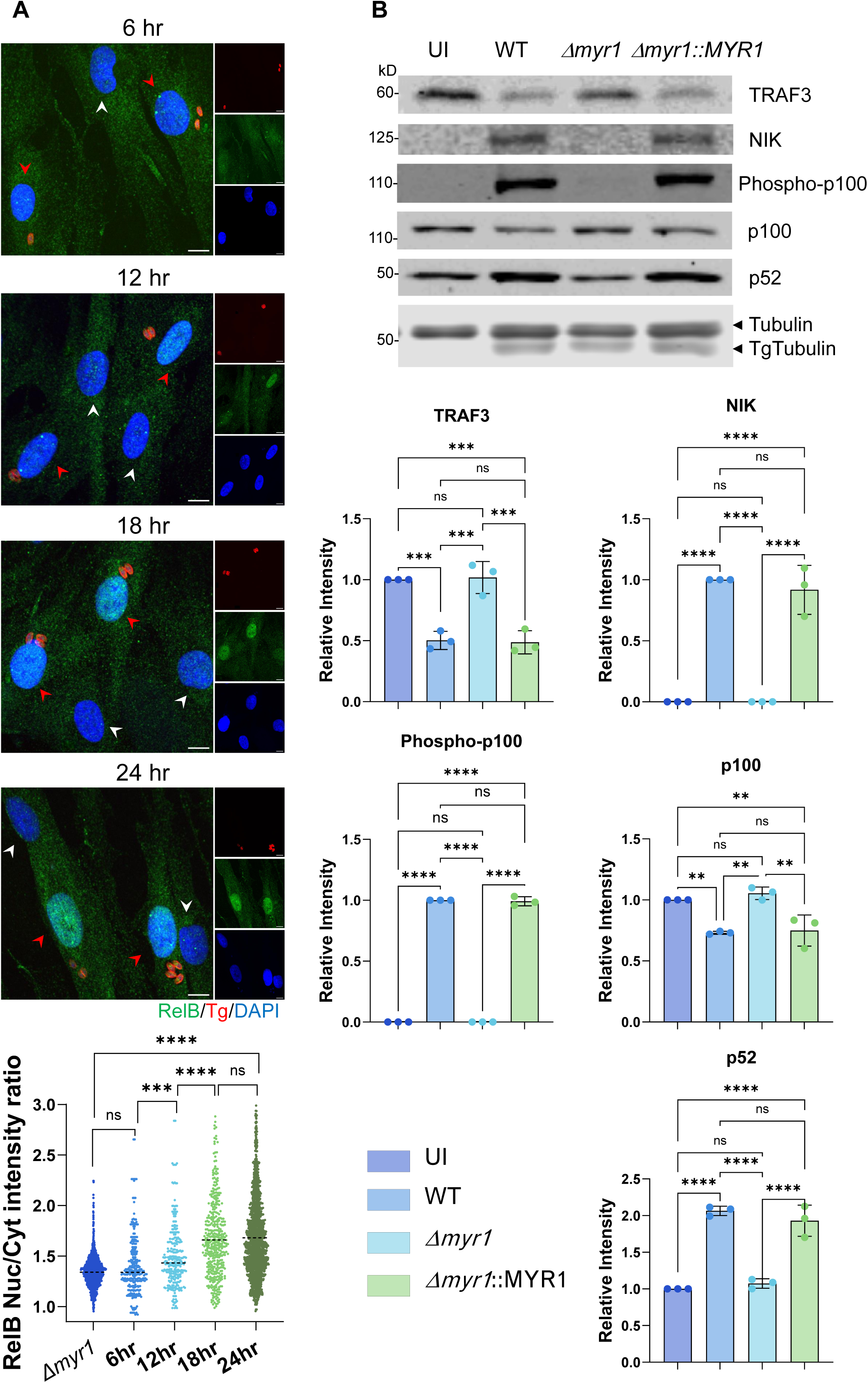
*T. gondii* activates the non-canonical NF-κB pathway through MYR1-dependent TRAF3 depletion and NIK stabilization. **(A)** Representative images (top) and quantitative analysis (bottom) for RelB nuclear accumulation in HFFs infected with RH (WT) parasites over a 24-h time course. At 6, 12, 18, and 24 h post-infection, cells were fixed and labeled with DAPI (blue), anti-GAP45 (red), and anti-RelB (green). Scale bars = 10 µm. Plots display the nuclear-to-cytoplasmic intensity ratios for at least 300 infected cells (red arrows) per condition; uninfected cells are indicated by white arrows. Data from three independent experiments were combined for analysis. The horizontal dashed lines indicate the mean. Statistical significance was determined using a one-way ANOVA with Tukey’s multiple comparison test; ***P < 0.001, ****P < 0.0001, ns = not significant. **(B)** Immunoblot analysis (top) and corresponding quantification (bottom) of TRAF3, NIK, p100 phosphorylation, and p100-to-p52 processing in HFFs that were uninfected (UI) or infected with RH (WT), *Δmyr1* or *Δmyr1*::MYR1 complement parasites. Lysates collected 24 h post-infection were resolved by SDS-PAGE and probed with specific primary antibodies; β-tubulin served as a loading control. Band intensities were measured using ImageJ and normalized to β-tubulin. Data are presented as mean ±SD from three biological replicates. Statistical significance for all panels was determined using a one-way ANOVA with Tukey’s multiple comparison test; **P < 0.01, ***P < 0.001, ****P < 0.0001, ns = not significant

To elucidate the mechanism by which *T. gondii* drives RelB/p52 activation, we examined upstream components of the non-canonical NF-κB pathway. In this pathway, TRAF3 constrains NIK abundance by promoting its constitutive degradation, and loss of TRAF3 stabilizes NIK to initiate downstream signaling (35, 41). Consistent with this mechanism, lysates from cells infected with WT or (Δ*myr1*::MYR1) complement parasites showed a marked reduction in TRAF3 protein levels compared to uninfected or Δ*myr1*-infected cells (Fig. 3B). This MYR1-dependent depletion of TRAF3 coincided with robust NIK stabilization, p100 phosphorylation, and increased p52 processing (Fig. 3B). Together, these results indicate that *T. gondii* engages the non-canonical NF-κB pathway, with MYR1-dependent effector export required for TRAF3 loss, NIK stabilization, and p100-to-p52 processing (Fig. 3B).

### The known MYR1-dependent effector set cooperates to drive nuclear RelB accumulation

To determine which MYR1-dependent effector(s) might drive this phenotype, we screened a panel of knockout parasites lacking individual secreted factors currently known to traffic to the host nucleus or cytoplasm. We tested single deletion mutants for GRA16, GRA18, GRA24, GRA28, GRA84, IST, NSM, and HCE1/TEEGR. Surprisingly, none of the single knockouts detectably reduced nuclear RelB accumulation, as all strains induced nuclear RelB at levels comparable to WT infection (Fig. 4A).

**Figure 4.**
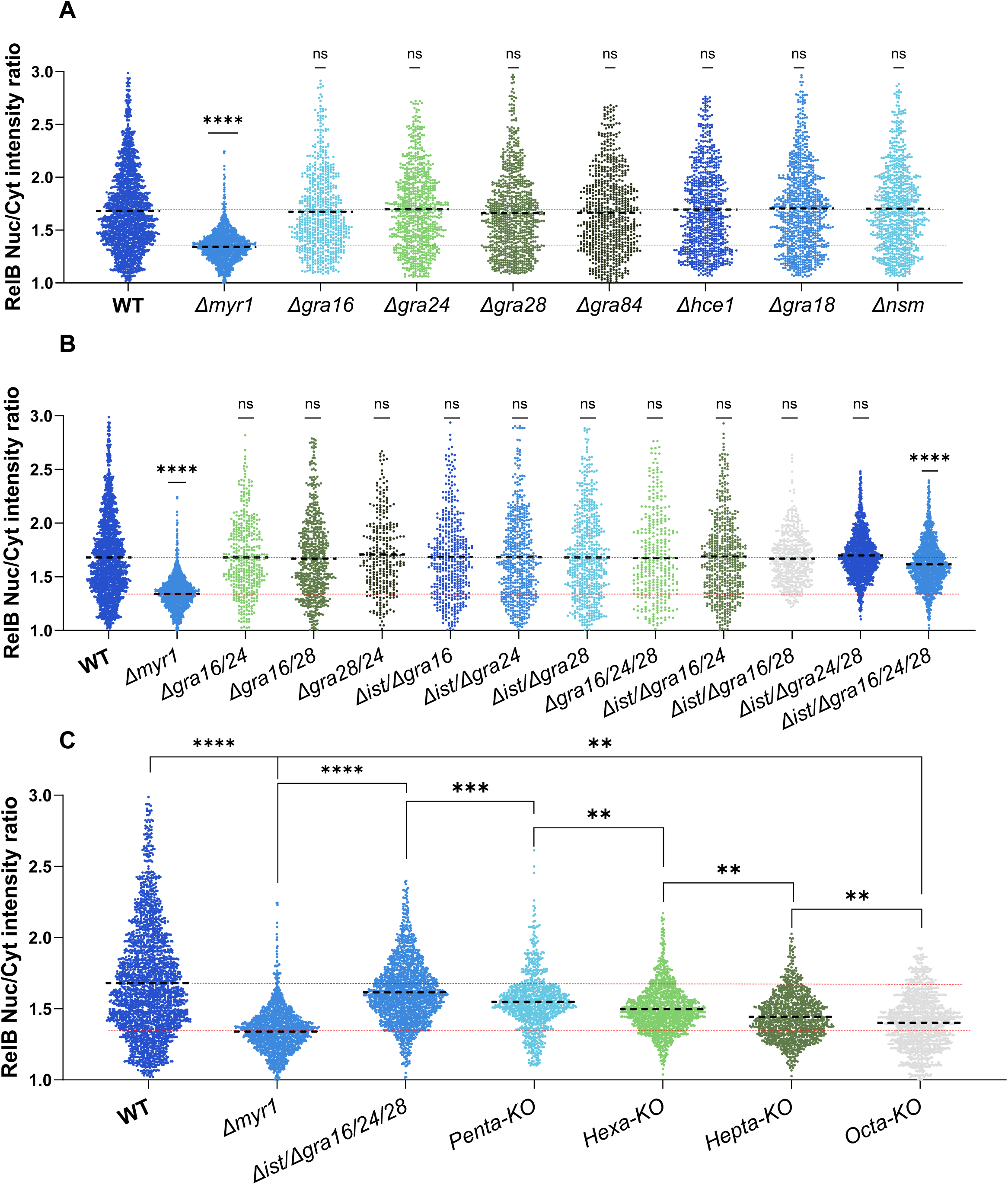
A suite of eight MYR1-dependent effectors cooperates to drive RelB nuclear accumulation. **(A)** Quantification of RelB nuclear accumulation in HFFs infected with single MYR1-dependent exported effector knockouts: *Δgra16, Δgra24, Δgra28, Δgra84, Δist, Δhce1, Δgra18, Δnsm*. **(B)** RelB nuclear-to-cytoplasmic intensity ratios in HFFs infected with various lower-order combinatorial knockout strains. **(C)** Progressive reduction of RelB nuclear accumulation in HFFs infected with higher-order combinatorial mutants: quadruple knockout (*Δist/Δgra16/Δgra24/Δgra28*), Penta-KO (*Δist/Δgra16/Δgra24/Δgra28/Δgra84*), Hexa-KO (*Δist/Δgra16/Δgra24/Δgra28/Δgra84/Δhce1*), Hepta-KO (*Δist/Δgra16/Δgra24/Δgra28/Δgra84/Δhce1/Δgra18*) and Octa-KO (*Δist/Δgra16/Δgra24/Δgra28/Δgra84/Δhce1/Δgra18/Δnsm*). **(A-C)** Plots display the nuclear-to-cytoplasmic intensity ratios for at least 300 infected cells per condition. The horizontal dashed lines indicate the mean. Data from three independent experiments were combined for analysis. Statistical significance was determined using a one-way ANOVA with Dunnett’s multiple comparison test; **P<0.01,***P < 0.001, ****P < 0.0001, ns = not significant.

The lack of a phenotype in single knockouts suggested substantial functional redundancy, consistent with emerging evidence that *T. gondii* effectors often act in overlapping combinations to shape host transcriptional outputs (25, 26). To address this, we leveraged a panel of combinatorial knockout parasites that we previously generated while dissecting MYR1-dependent suppression of IFN-γ-responsive transcription, focusing on IST, GRA16, GRA24, and GRA28 (25). Percent reductions for combinatorial strains were calculated relative to the dynamic range between the WT ratio (∼1.68) and the *Δmyr1* baseline (∼1.34). While single mutants and lower order combinations did not significantly reduce nuclear RelB accumulation, the quadruple knockout (*Δist/Δgra16/Δgra24/Δgra28*) lowered the nuclear-to-cytoplasmic ratio to ∼1.61, representing an approximately 20% shift toward the *Δmyr1* baseline (Fig. 4B). This partial reduction suggested the phenotype is distributed beyond just this core set of IFN-γ-influencing effectors. We therefore expanded the set using our IFN-γ CRISPR screen as a guide and prioritized GRA84, which ranked 209 among negatively selected genes under IFN-γ pressure (25). Deleting GRA84 in the quadruple background (*Δist/Δgra16/Δgra24/Δgra28/Δgra84*) strengthened the defect, yielding a ratio of ∼1.55, or an approximately 40% reduction in nuclear RelB relative to WT infection (Fig. 4C).

To further define the effector repertoire driving this phenotype, we expanded combinatorial mutagenesis to include additional MYR1-dependent effectors. We next deleted HCE1/TEEGR in the quintuple background, given its established role in modulating NF-κB-dependent transcription through EZH2 recruitment (18). The resulting hextuple mutant (*Δist/Δgra16/Δgra24/Δgra28/Δgra84/Δhce1*) further reduced the ratio to ∼1.49, an approximately 54% decrease relative to WT (Fig. 4C). We then targeted GRA18 (23). Although GRA18 is best known for stabilizing β-catenin, extensive crosstalk between Wnt and NF-κB signaling suggested it could contribute to this phenotype (23, 42). Deleting GRA18 in the hextuple background produced a cumulative ratio of ∼1.44, representing an approximately 70% reduction in nuclear RelB (Fig. 4C). Finally, to complete the analysis of the remaining candidate effector set, we deleted NSM in the septuple background. The resulting octuple knockout (*Δist/Δgra16/Δgra24/Δgra28/Δgra84/Δhce1/Δgra18/Δnsm)* exhibited a ratio of ∼1.4, an approximately 80% reduction compared to WT infection (Fig. 4C). Collectively, these data confirm substantial functional redundancy and indicate that this suite of eight MYR1-dependent effectors cooperates to drive most of the non-canonical NF-κB response during *T. gondii* infection.

## Discussion

The co-evolutionary struggle between *T. gondii* and the host immune system is perhaps most clearly defined by the parasite’s sophisticated manipulation of the NF-κB signaling axis. While much of our understanding has centered on the subversion of canonical p65/p50 signaling, our findings identify activation of the non-canonical NF-κB pathway as a previously underappreciated outcome of the tachyzoite stage of the infection. We show that *T. gondii* induces robust nuclear accumulation of RelB and p52 across both Type I and Type II genetic backgrounds in human and mouse fibroblasts. Importantly, this host cell reprogramming is strictly dependent on MYR1-mediated effector export, identifying dense granule translocation as the proximal determinant of the non-canonical response.

Our analysis of upstream pathway structure suggests that *T. gondii* engages non-canonical NF-κB at the level of the NIK regulatory checkpoint. WT infection triggers depletion of TRAF3, stabilization of NIK, and subsequent phosphorylation and processing of p100, all of which are abolished in the absence of MYR1. This model is strongly supported by the established architecture of non-canonical NF-κB signaling, in which TRAF3 serves as a key negative regulator of NIK abundance and p100 processing to p52, thereby licensing RelB/p52 nuclear signaling (35, 43). Mechanistically, the MYR1-dependent effector suite implicated here is composed of established transcriptional modulators that reprogram host gene expression, suggesting that *T. gondii* drives non-canonical NF-κB activation by tuning the expression and stoichiometry of checkpoint components, rather than by a single direct cytoplasmic interaction (44). For example, coordinated changes in the abundance of TRAF3, TRAF2, or cIAP1/2 could lower the threshold for NIK accumulation and favor sustained signaling once initiated (33, 35). This mode of control is also consistent with the gradual, persistent kinetics of RelB nuclear accumulation observed over 24 hours (40).

The observation that non-canonical RelB/p52 activation is conserved across both Type I and Type II strains, while canonical RelA/p65 activation is a unique feature of Type II infection, suggests a tiered strategy for host NF-κB reprogramming (28, 29). In this model, the engagement of the non-canonical pathway could represent a baseline component of host cell subversion, driven by a distributed network of eight MYR1-dependent effectors common to both lineages. In contrast, the GRA15-mediated activation of RelA/p65 in Type II strains overlays a specialized inflammatory signature onto this basal state (29). One potential consequence of this dual activation is the formation of RelA-RelB heterodimers, which have been reported to possess distinct DNA-binding properties and to participate in regulatory feedback that can temper acute inflammatory responses (45). By maintaining a conserved non-canonical response across lineages while varying the canonical input, the parasite may differentially shape the host signaling equilibrium, potentially balancing host cell viability with the evasion of sterile immunity across diverse genetic backgrounds.

A central conceptual finding is that non-canonical NF-κB activation is not attributable to a single exported effector. Across a panel of parasites lacking individual MYR1-dependent effectors, none of the single deletions significantly diminished nuclear RelB accumulation. Instead, the phenotype emerged through a cumulative, multi-effector regulatory network, with progressively larger defects observed as additional effectors were deleted in higher-order mutant backgrounds. This progressive reduction suggests that the eight effectors identified here: GRA16, GRA18, GRA24, GRA28, GRA84, IST, NSM, and HCE1/TEEGR do not merely provide redundancy in the classical sense, but rather function through additive cooperation to reach a signaling threshold. This distributed logic mirrors the strategies described for the suppression of IFN-γ responsive transcription and the tuning of canonical NF-κB outputs, extending this principle to pathway choice within the NF-κB system itself (18, 25, 26).

Understanding why *T. gondii* invests so heavily in multi-node targeting benefits from considering the host’s own architecture. Innate signaling and transcriptional circuits are characterized by profound transcriptional robustness, shaped by evolution to maintain stable outputs despite genetic variation and environmental noise (46, 47). Population-scale human genome analyses support this robustness, identifying predicted biallelic loss-of-function genotypes in a subset of NF-κB pathway genes, including core components such as p65 and IKKα, as well as regulators linked to non-canonical pathway control such as ZFP91 and TRIM55 (48–51). Although some of these genotypes may have incompletely characterized clinical consequences or present as specific immune phenotypes rather than being uniformly benign, their presence in ostensibly healthy populations underscores the inherent plasticity of the host signaling network. This stability is maintained through multiple layers of control, including functional redundancy between transcription factor paralogs and the existence of parallel signaling cascades (46).

Consistent with this idea, NF-κB upstream signaling modules include seemingly redundant components in which loss of one family member can be partially absorbed by others, as illustrated by partial compensation of IKKβ loss by IKKα in vivo (52). In such a stable system, a parasite strategy that relies on a single effector to trigger a specific molecular change would likely be buffered and neutralized by the host’s internal backups (33, 53). By deploying a suite of effectors with overlapping activities, *T. gondii* can push multiple control points simultaneously to successfully and appropriately modulate this host function, converting a stable circuit into one that is permissive for sustained non-canonical signaling.

Engaging non-canonical NF-κB during infection may reflect a trade-off between host defense and parasite persistence. While RelB is required for protective IFN-γ responses in vivo (36), non-canonical NF-κB signaling is also associated with slower, homeostatic transcriptional programs and enhanced cell survival (33, 54). RelB/p52 complexes regulate a subset of anti-apoptotic genes, including Bcl-xL (BCL2L1), Bfl-1/A1 (BCL2A1), and c-IAP2 (BIRC3), and can promote cellular states distinct from the acute inflammatory outputs driven by p65 (55–57). It is therefore plausible that, in non-hematopoietic cells such as fibroblasts, controlled activation of RelB favors host cell survival and stabilization of the intracellular niche. In this model, RelB activation would not simply amplify host resistance, but instead reshape transcriptional programs in a manner that balances inflammatory signaling with cellular viability, potentially benefiting parasite persistence. This idea is consistent with our prior observation that MYR1-deficient parasites replicate less and egress earlier than wild-type under unstimulated conditions, suggesting that MYR1-dependent effector export helps sustain the intracellular niche (25).

Several important questions follow from the results reported here. First, defining the RelB-dependent transcriptional program specifically induced during *T. gondii* infection will be essential to determine whether parasite-driven RelB activation reinforces, diverts, or partially rewires canonical immune outputs. Second, although higher-order mutants substantially reduce nuclear RelB accumulation, a residual signal remains, raising the possibility that additional, yet-to-be-discovered exported factors or indirect host-mediated inputs contribute to pathway engagement. Future experiments that pair RelB loss of function with transcriptomic profiling and phenotypic readouts will be essential to clarify how this axis shapes the balance between parasite fitness and host cell fate.

## Materials and Methods

### Parasite and host cell culture

*T. gondii* tachyzoites were serially passaged in human foreskin fibroblast (HFF) monolayers cultured in D10 medium (Dulbecco’s modified Eagle’s medium [DMEM; Invitrogen]) supplemented with 10% HyClone Cosmic calf serum (Cytiva), 10 µg/mL gentamicin (Thermo Fisher Scientific), 2 mM glutamine (Thermo Fisher Scientific). MEF cells were maintained in D10. All strains and host cell lines were determined to be mycoplasma-negative using the e-Myco Plus Kit (Intron Biotechnology). Strains used in this study are listed in Table S1. Gene disruptant lines were generated using CRISPR/Cas9 (58), as described below. Parasite lines generated in other studies include RH*ΔhxgprtΔku80* (59), ME49 *FLuc* (60), RH *Δgra84* (22).

### Plasmid construction and genome editing

Plasmids were generated by site-directed mutagenesis of existing plasmids or assembled from DNA fragments by the Gibson method (61). All plasmids used in this study are listed in Table S2.

### Primers

All primers were synthesized by Sigma. All CRISPR/Cas9 plasmids used in this study were derived from the single-guide RNA (sgRNA) plasmid pSAG1:CAS9-GFP, U6:sgUPRT (58) by Q5 site-directed mutagenesis (New England Biolabs) to alter the 20-nt sgRNA sequence, as described previously (62). Primers for plasmids are listed in Table S3.

### Parasite transfection

Following natural egress, freshly harvested extracellular parasites were transfected with plasmids as previously described (58). Briefly, approximately 2 × 10^7 parasites were resuspended in 370 µL cytomix buffer and mixed with approximately 30 µL purified plasmid or amplicon DNA in a 4 mm gap BTX cuvette. Parasites were electroporated using a BTX ECM 830 electroporator (Harvard Apparatus) with the following settings: 1,700 V, 176 µs pulse length, 2 pulses, and a 100 ms interval between pulses. Transgenic parasites were selected by outgrowth under drug selection as needed using mycophenolic acid (25 µg/mL) plus xanthine (50 µg/mL) (MPA/Xa), pyrimethamine (3 µM), chloramphenicol (40 µM), 5-fluorodeoxyuridine (10 µM) (Sigma), or phleomycin (5 µg/mL) (InvivoGen). Stable clones were isolated by limiting dilution on HFF monolayers in 96-well plates.

### Generation of quintuple, sextuple, septuple, and octuple knockout strains

Because available selectable markers were limiting, 5-through 8-gene MYR1-dependent knockout strains were generated by fluorescence sorting and subcloning without drug selection. Parasites were transfected with the relevant guide RNA(s) together with an amplicon donor DNA. Donor amplicons were generated using genomic DNA from the appropriate single knockout strain as template, amplifying the selectable cassette together with approximately 500 bp of upstream and 500 bp of downstream homology arms flanking the cassette. Guide RNA(s) and donor amplicon were mixed at a 1:3 ratio to increase the likelihood that each Cas9-GFP positive parasite also received donor DNA (Fig. S3).

### Immunofluorescence assay (IFA) and image analysis

Host cells were seeded onto glass coverslips in 24-well plates and cultured for 24 h prior to infection. Cells were infected the following day with *T. gondii* (1 × 10^5 parasites per well). At 24 h post-infection, infected cells were fixed with 4% formaldehyde for 10 min at room temperature (RT), washed with phosphate-buffered saline (PBS), permeabilized with 0.2% Triton X-100 for 10 min, and blocked in PBS containing 10% normal goat serum (Thermo) (PBS with 10% NGS).

Coverslips were washed four times in PBS (5 min per wash) and incubated with primary antibodies diluted in blocking buffer overnight at 4°C. The following primary antibodies were used: rabbit anti-RelB (1:300; Proteintech, 25027-1-AP), mouse anti-NFκB p52 (1:100; Santa Cruz Biotechnology, sc-7386), and mouse anti-GAP45 or rabbit anti-GAP45 (kindly provided by Dr D. Etheridge, University of Georgia, USA). Samples were washed four times in PBS (5 min per wash) and incubated with secondary antibodies diluted in blocking buffer containing Hoechst 33342 (1 µg/mL). Secondary antibodies were goat anti-rabbit Alexa Fluor 488 (1:2,000) and goat anti-mouse Alexa Fluor 488 (1:2,000), (Invitrogen). *T. gondii* containing vacuoles were stained with anti-GAP45 and visualized with Alexa Fluor 568 (Invitrogen). Following secondary staining, coverslips were washed four times in PBS (5 min per wash) and mounted on slides using ProLong Gold Antifade Reagent (Invitrogen).

Samples were imaged using a Plan-Apochromat 20× objective (Zeiss) on a 3i Marianas Plus CSU-W1 SoRa super-resolution spinning disk confocal microscope equipped with a LaserStack (3i) laser source and a Prime 95B CMOS digital camera (Teledyne Photometrics). Images were acquired using SlideBook software.

Raw immunofluorescence images were exported from SlideBook as single-channel 16-bit TIFF files and analyzed in CellProfiler (version 4.2.8). For each field of view, matched 3-channel image sets (DAPI, GAP45, and RelB or p52) were processed together. Host nuclei were segmented from the DAPI channel using size-based filtering to exclude debris and merged nuclei. Parasites were segmented independently from the GAP45 channel using size-based gating. Whole-cell regions were approximated by expanding outward from each nucleus using the RelB or p52 signal, which is distributed throughout the host cell. Cytoplasmic regions were then defined by subtracting the nuclear mask from the whole-cell mask. Parasites were assigned to host cells to classify cells as infected or uninfected, and this classification was propagated to the corresponding nuclei. Parasite regions were excluded from cytoplasmic measurements to avoid contamination of host fluorescence measurements by parasite-derived signal.

RelB or p52 fluorescence was quantified as mean pixel intensity within the nuclear and cytoplasmic compartments, and a nuclear-to-cytoplasmic mean intensity ratio was calculated for each cell. Nuclear-to-cytoplasmic ratios were quantified in at least 100 infected cells per group in each biological replicate across at least three independent experiments. Measurement outputs were exported as spreadsheets, organized in Microsoft Excel, and analyzed and plotted in GraphPad Prism.

### Immunoblotting

HFFs grown in T-12.5 flasks were infected with 5 × 10^6 parasites per flask. At 24 h post-infection, monolayers were washed with PBS, scraped into ice-cold PBS, and pelleted at 500 × g. Cell pellets were lysed in RIPA buffer (50 mM Tris-HCl, pH 7.4; 150 mM NaCl; 1% Triton X-100; 0.5% sodium deoxycholate; 0.1% SDS) supplemented with 1× Halt protease and phosphatase inhibitor cocktail.

Protein samples were prepared in Laemmli buffer containing 100 mM dithiothreitol, boiled for 5 min, resolved by SDS-PAGE on 10% polyacrylamide gels, and transferred to nitrocellulose membranes. Membranes were blocked in PBS containing 2.5% fat-free milk, then incubated with primary antibodies diluted in blocking buffer containing 0.1% Tween 20. Membranes were washed in PBS with 0.1% Tween 20 and incubated with goat IRDye-conjugated secondary antibodies (LI-COR Biosciences) diluted in blocking buffer (as indicated in the figure legends). After additional washes, membranes were scanned on a LI-COR Odyssey imaging system (LI-COR Biosciences). Band intensities were quantified using ImageJ.

Primary antibodies used were: rabbit anti-phospho-NF-kappaB2 p100 (Ser866/870) (1:1,000; Cell Signaling Technology, #4810), rabbit anti-NF-kappaB2 p100/p52 (1:1,000; Cell Signaling Technology, #4882), rabbit anti-NIK (1:1,000; Cell Signaling Technology, #4994), rabbit anti-TRAF3 (1:1,000; Cell Signaling Technology, #4729), and mouse anti-β-tubulin (Developmental Studies Hybridoma Bank, AB_528499), which detects both parasite and host β-tubulin and was used as a loading control.

### Statistical Analysis

Statistical analysis for each experiment was performed on the combined data in PRISM (GraphPad). Replicates, significance values, and tests performed are included in the figure legends.

## Acknowledgments

We thank Drew Etheridge lab for help and reagent support; Juan Bustamante at the CTEGD Cytometry Shared Resource Lab for technical assistance.

This study was supported by NIH grant (RO1 AI129529) to J.B. and UGA startup fund to A.R.

Conceptualization, A.R., M.P. and J.C.B.; methodology, A.R. and M.P.; investigation, K.B., M.P., A.R.; formal analysis, K.B., M.P., A.R.; writing-original draft, A.R.; writing-review & editing, K.B., A.R, M.B. and J.C.B.; funding acquisition, J.C.B. and A.R.; resources, S.K, A.R. and J.C.B.; supervision, A.R. and J.C.B.

## Conflicts

The authors declare no conflicts.

## Supplemental items

**Table S1.** *T. gondii* strains used in this study.

**Table S2.** Plasmids used in this study.

**Table S3.** Primers used in this study.

---

**Figure S1.**
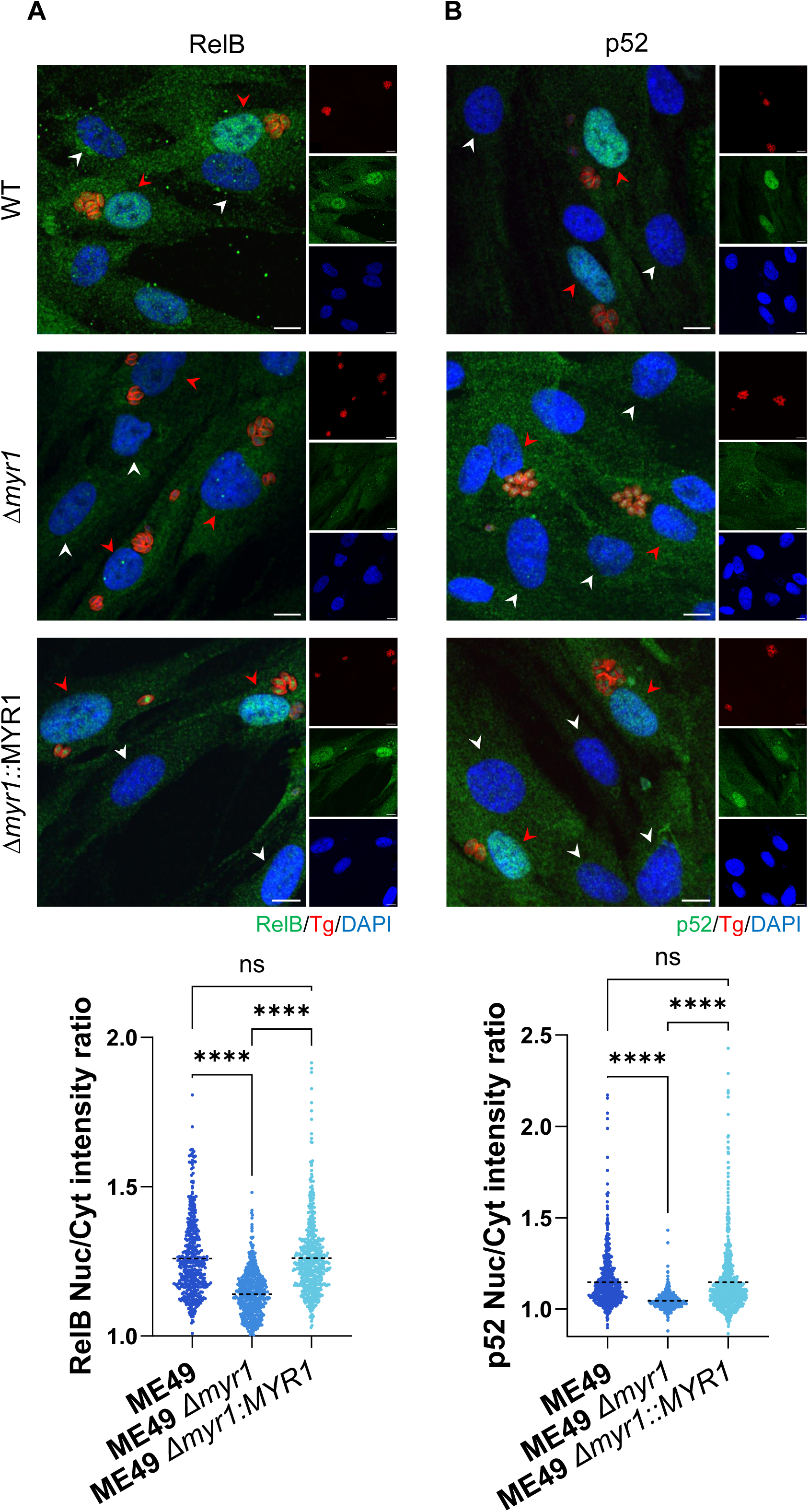
Conservation of MYR1-dependent non-canonical NF-κB activation in Type II (ME49) parasites. A-B) Representative images (top) and quantitative analysis (bottom) for RelB **(A)** and p52 **(B)** nuclear accumulation in HFFs infected with Type II ME49 (WT), Δ*myr1* or Δ*myr1*::MYR1 complement parasites. Twenty-four hours post-infection, cells were fixed and labeled with DAPI (blue), anti-GAP45 (red), and either anti-RelB or anti-p52 (green). Scale bars = 10 µm. Plots display the nuclear-to-cytoplasmic signal ratios for at least 300 infected cells (red arrows) per condition; uninfected cells are indicated by white arrows. Data from three independent experiments were combined for analysis. The horizontal dashed lines indicate the mean. Statistical significance was determined using a one-way ANOVA with Tukey’s multiple comparison test; ****P < 0.0001, ns = not significant.

**Figure S2.**
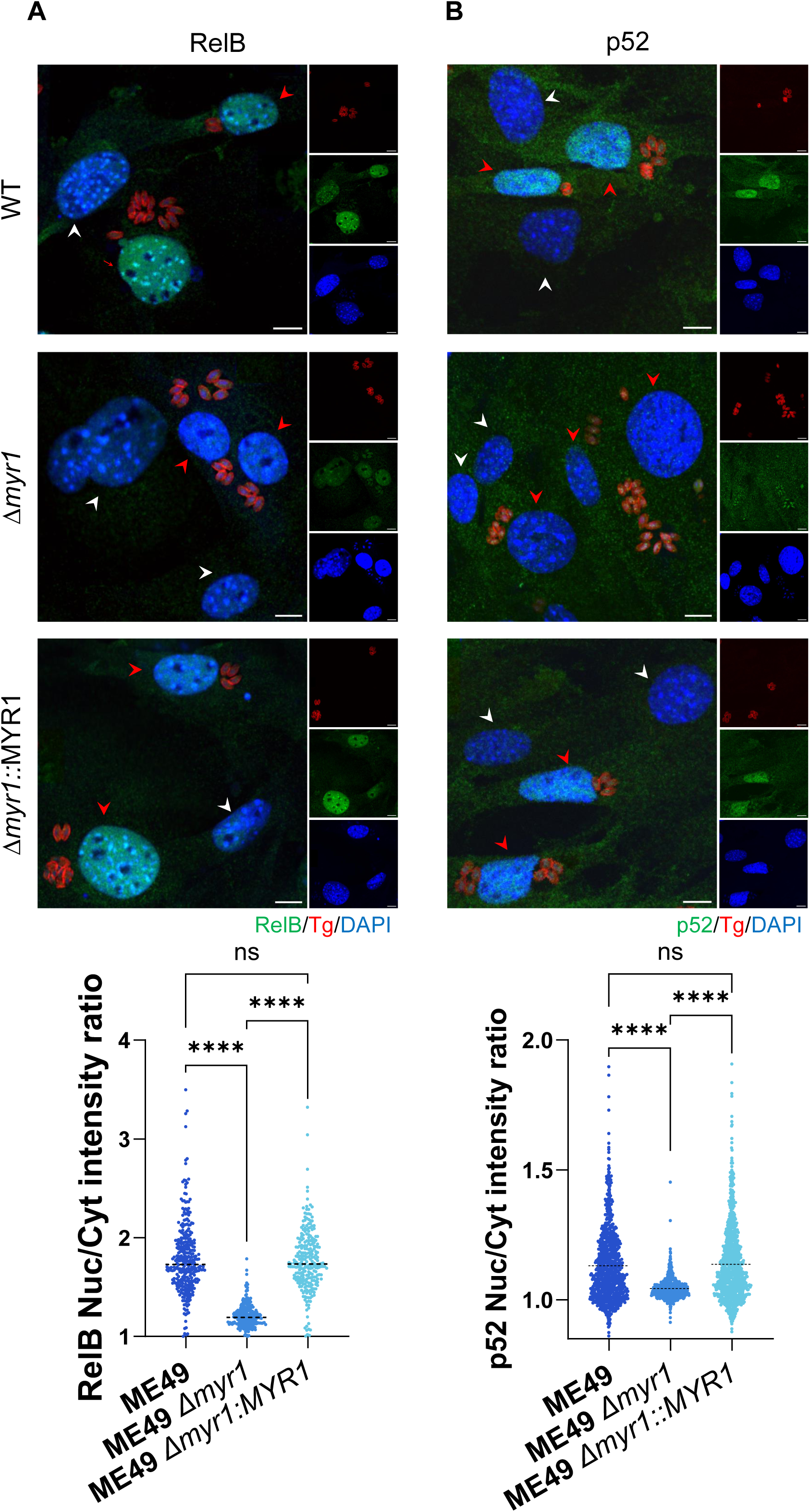
Type II (ME49) *T. gondii* induces MYR1-dependent RelB and p52 accumulation in murine cells. **A-B)** Representative images (top) and quantitative analysis (bottom) for RelB **(A)** and p52 **(B)** nuclear accumulation in MEFs infected with Type II ME49 (WT), Δ*myr1* or Δ*myr1*::MYR1 complement parasites. Twenty-four hours post-infection, cells were fixed and labeled with DAPI (blue), anti-GAP45 (red), and either anti-RelB or anti-p52 (green). Scale bars = 10 µm. Plots display the nuclear-to-cytoplasmic signal ratios for at least 300 infected cells (red arrows) per condition; uninfected cells are indicated by white arrows. Data from three independent experiments were combined for analysis. Statistical significance was determined using a one-way ANOVA with Tukey’s multiple comparison test; ****P < 0.0001, ns = not significant.

**Figure S3.**
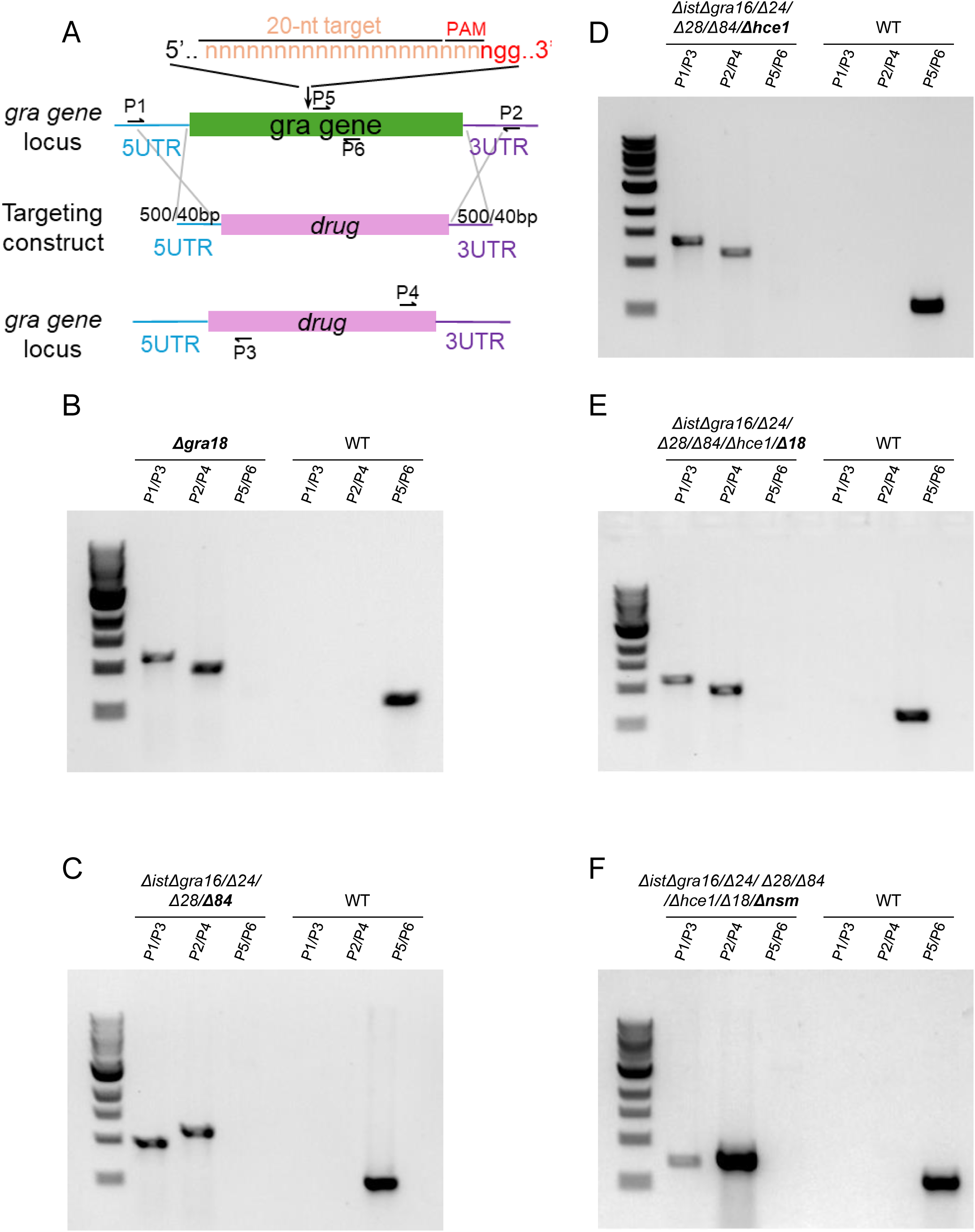
Generation and molecular validation of combinatorial knockout parasite strains. **(A)** Schematic representation of the CRISPR/Cas9-mediated gene deletion strategy used to generate the transgenic strains in this study. Single sgRNA-expressing CRISPR/Cas9 (gRNA) plasmids targeting the middle of each gene were used to introduce a double-strand break and facilitate isolation of knockouts by homologous recombination. For single mutants, targeting constructs consisted of a selectable cassette and short homology flanks (∼40 bp) positioned immediately upstream of the translation initiation site (left arm) and downstream of the stop codon (right arm). For penta-through octa-KO strains, donor amplicons containing ∼500 bp homology arms flanking the selectable cassette were used, followed by fluorescence sorting and subcloning without drug selection. (**B-F**) Diagnostic PCR verification of locus disruption for the indicated mutant strains: (B) Δgra18; (C) penta-KO, generated by addition of Δgra84 to the tetra-KO background (*Δist/Δgra16/Δgra24/Δgra28/Δgra84*); (D) hexa-KO, generated by addition of Δhce1 to the penta-KO background (*Δist/Δgra16/Δgra24/Δgra28/Δgra84/Δhce1*); (E) hepta-KO, generated by addition of Δgra18 to the hexa-KO background (*Δist/Δgra16/Δgra24/Δgra28/Δgra84/Δhce1/Δgra18*); and (F) octa-KO, generated by addition of Δnsm to the hepta-KO background (*Δist/Δgra16/Δgra24/Δgra28/Δgra84/Δhce1/Δgra18/Δnsm*). Primer binding sites are indicated in (**A**): P1/P3 and P2/P4 confirm integration of the left and right homology arms, respectively; P5/P6 examines the integrity of the endogenous locus. Successful knockout clones yielded PCR products with P1/P3 and P2/P4 but no product with P5/P6, whereas wild-type parasites showed the opposite pattern. See Table S3 for oligonucleotide sequences.

## Notes

### Competing Interest Statement

The authors have declared no competing interest.

### Summary of Updates

Supplemental figures have been included in the manuscript.

